# Gene model for the ortholog of *Glys in Drosophila ananassae*

**DOI:** 10.1101/2025.07.09.662856

**Authors:** Madeline L. Gruys, Abigail R. Myers, Cole A. Kiser, Leon F. Laskowski, Isaac Romo, Amber Marie Richardson, James J. Youngblom, Chinmay P. Rele, Laura K. Reed

## Abstract

Gene model for the ortholog of *Glycogen synthase* (*Glys*) in the May 2011 (Agencourt dana_caf1/DanaCAF1) Genome Assembly (GenBank Accession: GCA_000005115.1) of *Drosophila ananassae*. This ortholog was characterized as part of a developing dataset to study the evolution of the Insulin/insulin-like growth factor signaling pathway (IIS) across the genus *Drosophila* using the Genomics Education Partnership gene annotation protocol for Course-based Undergraduate Research Experiences.

## Introduction

*This article reports a predicted gene model generated by undergraduate work using a structured gene model annotation protocol defined by the Genomics Education Partnership (GEP; thegep.org) for Course-based Undergraduate Research Experience (CURE). The following information in quotes may be repeated in other articles submitted by participants using the same GEP CURE protocol for annotating Drosophila species orthologs of Drosophila melanogaster genes in the insulin signaling pathway*.

“In this GEP CURE protocol students use web-based tools to manually annotate genes in non-model *Drosophila* species based on orthology to genes in the well-annotated model organism fruitfly *Drosophila melanogaster*. The GEP uses web-based tools to allow undergraduates to participate in course-based research by generating manual annotations of genes in non-model species (Rele et al., 2023). Computational-based gene predictions in any organism are often improved by careful manual annotation and curation, allowing for more accurate analyses of gene and genome evolution (Mudge and Harrow 2016; Tello-Ruiz et al., 2019). These models of orthologous genes across species, such as the one presented here, then provide a reliable basis for further evolutionary genomic analyses when made available to the scientific community.” (Myers et al., 2024).

“The particular gene ortholog described here was characterized as part of a developing dataset to study the evolution of the Insulin/insulin-like growth factor signaling pathway (IIS) across the genus *Drosophila*. The Insulin/insulin-like growth factor signaling pathway (IIS) is a highly conserved signaling pathway in animals and is central to mediating organismal responses to nutrients (Hietakangas and Cohen 2009; Grewal 2009).” (Myers et al., 2024).

“*Glycogen synthase* (*Glys*; aka. *GS*) is a gene within the Insulin-signaling pathway in *Drosophila* and encodes a glycosyltransferase that catalyzes linkage of glucose monomers into glycogen. Glys activity is regulated allosterically by glucose 6-phosphate and phosphorylation/dephosphorylation allowing for control of cellular glycogen levels (Plyte et al., 1992; Roach et al., 2012). Null *Glys* mutants exhibit growth defects and reduced larval viability in *Drosophila* (Yamada et al., 2019).” (Backlund et al., 2025).

“*D. ananassae* (NCBI:txid7217) is part of the *melanogaster* species group within the subgenus *Sophophora* of the genus *Drosophila* (Sturtevant 1939; Bock and Wheeler 1972). It was first described by Doeschall (1858). *D. ananassae* is circumtropical (Markow and O’Grady 2005<u>; https://www.taxodros.uzh.ch</u>, accessed 1 Feb 2023), and often associated with human settlement (Singh 2010). It has been extensively studied as a model for its cytogenetic and genetic characteristics, and in experimental evolution (Kikkawa 1938; Singh and Yadav 2015).” (Lawson et al., 2024; Gruys et al., 2025a).

We propose a gene model for the *D. ananassae* ortholog of the *D. melanogaster Glycogen synthase* (*Glys*) gene. The genomic region of the ortholog corresponds to the uncharacterized protein LOC26515236 (RefSeq accession XP_032308811.1) in the May 2011 (Agencourt dana_caf1/DanaCAF1) Genome Assembly of *D. ananassae* (GenBank Accession: GCA_000005115.1; Drosophila 12 Genomes Consortium, 2007). This model is based on RNA-Seq data from *D. ananassae* (SRP006203, SRP007906; PRJNA257286, PRJNA388952; Brown et al., 2014; Rogers et al., 2014; Yang et al., 2018) and *Glys* in *D. melanogaster* using FlyBase release FB2024_02 (GCA_000001215.4; Gramates et al., 2022; Jenkins et al., 2022; Larkin et al., 2021).

### Synteny

The reference gene, *Glys*, occurs on chromosome 3R in *D. melanogaster* and is flanked upstream by *Guanylyl cyclase at 88E* (*Gyc88E*) and *Myofilin* (*Mf*) and downstream by *CG3987* and *CG3984*. The *tblastn* search of *D. melanogaster* Glys-PA (query) against the *D. ananassae* (GenBank Accession: GCA_000005115.1) Genome Assembly (database) placed the putative ortholog of *Glys* within scaffold_13340 (CH902617.1) at locus LOC26515236 (XP_032308811.1) — with an E-value of 0.0 and a percent identity of 86.86%. Furthermore, the putative ortholog is flanked upstream by LOC6500618 (XP_044570880.1) and LOC6499946 (XP_001953578.2), which correspond to *Gyc88E* and *Mf* in *D. melanogaster* (E-value: 0.0 and 0.0; identity: 85.35% and 94.05%, respectively, as determined by *blastp*; Figure 1A, Altschul et al., 1990). The putative ortholog of *Glys* is flanked downstream by LOC6500617 (XP_032308829.1) and LOC6500614 (XP_032308765.1), which correspond to *CG3987* and *CG3984* in *D. melanogaster* (E-value: 4e-144 and 6e-100; identity: 55.19% and 60.32%, respectively, as determined by *blastp*). The putative ortholog assignment for *Glys* in *D. ananassae* is supported by the following evidence: The genes surrounding the *Glys* ortholog are orthologous to the genes at the same locus in *D. melanogaster* and local synteny is completely conserved, supported by results generated from *blastp*; we conclude that LOC26515236 is the correct ortholog of *Glys* in *D. ananasssae* (Figure 1A).

**Figure 1:**
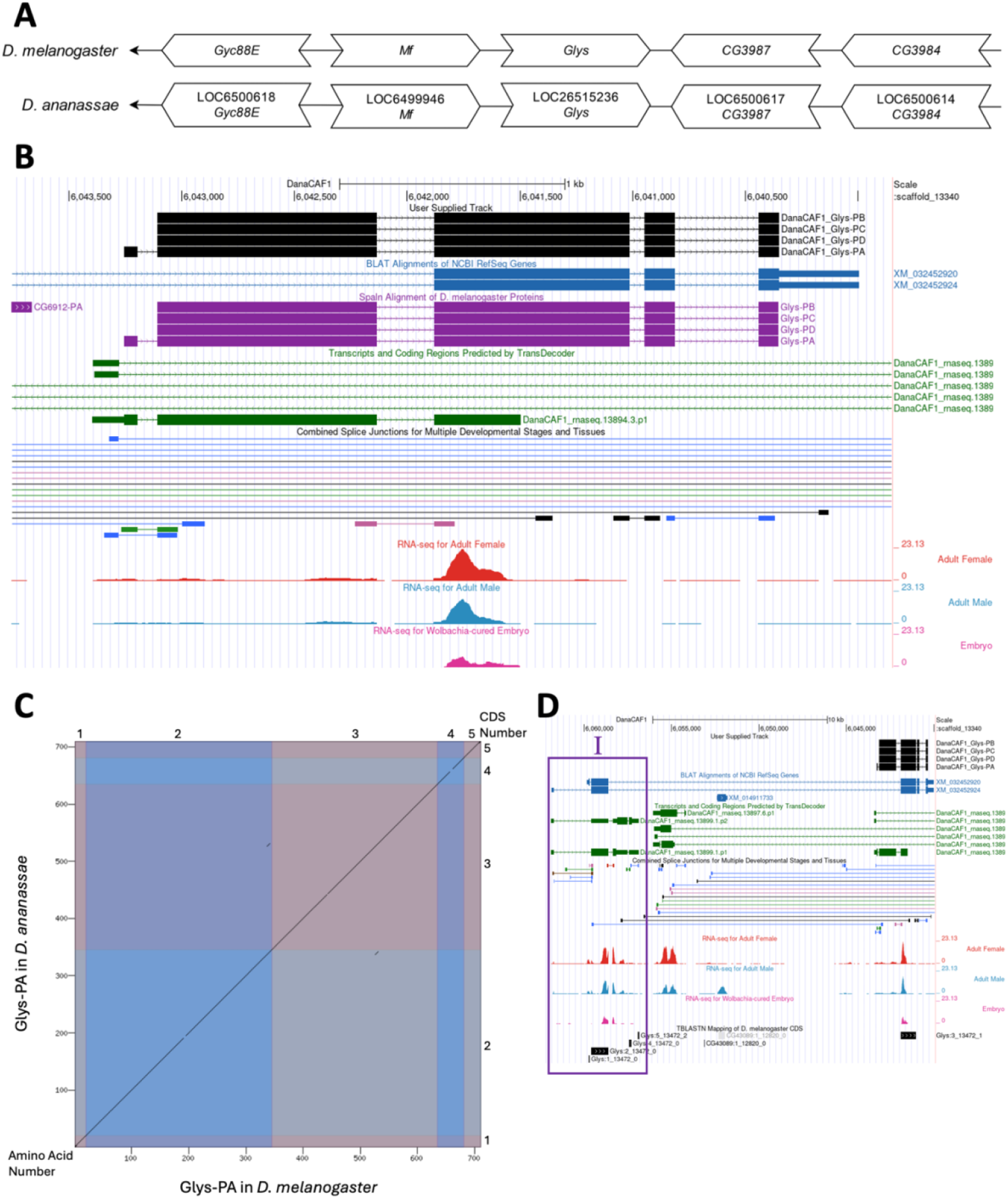
Glys gene model comparison between *Drosophila ananassae* and *Drosophila melanogaster* orthologs. **(A) Synteny comparison of the genomic neighborhoods for *Glys* in *Drosophila melanogaster* and *D. ananassae***. Thin underlying arrows indicate the DNA strand within which the reference gene–*Glys*–is located in *D. melanogaster* (top) and *D. ananassae* (bottom). Thin arrows pointing to the left indicate that *Glys* is on the negative (-) strand in *D. ananassae* and *D. melanogaster*. The wide gene arrows pointing in the same direction as *Glys* are on the same strand relative to the thin underlying arrows, while wide gene arrows pointing in the opposite direction of *Glys* are on the opposite strand relative to the thin underlying arrows. White gene arrows in *D. ananassae* indicate orthology to the corresponding gene in *D. melanogaster*. Gene symbols given in the *D. ananassae* gene arrows indicate the orthologous gene in *D. melanogaster*, while the locus identifiers are specific to D. *ananassae*. **(B) Gene Model in GEP UCSC Track Data Hub (Raney et al**., **2014)**. The coding-regions of *Glys* in *D. ananassae* are displayed in the User Supplied Track (black); coding CDSs are depicted by thick rectangles and introns by thin lines with arrows indicating the direction of transcription. Subsequent evidence tracks include *BLAT* Alignments of NCBI RefSeq Genes (dark blue, alignment of Ref-Seq genes for *D. ananassae*), *Spaln* of *D. melanogaster* Proteins (purple, alignment of Ref-Seq proteins from *D. melanogaster*), Transcripts and Coding Regions Predicted by *TransDecoder* (dark green), RNA-Seq from Adult Females, Adult Males, and Wolbachia-cured Embryos (red, light blue and pink, respectively; alignment of Illumina RNA-Seq reads from *D. ananassae*), and Splice Junctions Predicted by *regtools* using *D. ananassae* RNA-Seq (SRP006203, SRP007906; PRJNA257286, PRJNA388952). The splice junctions pertaining to the *Glys* ortholog, JUNC00120677 (black), JUNC00120692 and JUNC00120675 (blue), JUNC00120691 (green), and JUNC00120683 (pink) have read-depths of 7, 20, 28, 57 and 381, respectively. **(C) Dot Plot of Glys-PA in *D. melanogaster* (x-axis) vs. the orthologous peptide in *D. ananassae* (y-axis)**. Amino acid number is indicated along the left and bottom; CDS number is indicated along the top and right, and CDSs are also highlighted with alternating colors. Line breaks in the dot plot indicate mismatching amino acids at the specified location between species. **(D) Upstream region of *Glys* ortholog gene model displayed in *D. ananassae* UCSC Genome Browser**. The coding-regions of *Glys* in *D. ananassae* are displayed in the User Supplied Track (black); coding CDSs are depicted by thick rectangles and introns by thin lines with arrows indicating the direction of transcription. Subsequent evidence tracks include *BLAT* Alignments of NCBI RefSeq Genes (dark blue, alignment of Ref-Seq genes for *D. ananassae*), Transcripts and Coding Regions Predicted by *TransDecoder* (dark green), RNA-Seq from Adult Females, Adult Males, and Wolbachia-cured Embryos (red, light blue and pink, respectively; alignment of Illumina RNA-Seq reads from *D. ananassae*), Splice Junctions Predicted by *regtools* using *D. ananassae* RNA-Seq (SRP006203, SRP007906; PRJNA257286, PRJNA388952) and *tblastn* Mapping of *D. melanogaster* CDS (black and light grey). Splice junctions shown have a read-depth of 10-49, 50-99, 100-499, 500-999, >1000 supporting reads in blue, green, pink, brown, and red, respectively. The *BLAT* Alignments of NCBI RefSeq gene prediction that includes the putative ortholog in *D. ananassae* appears to exhibit shifted CDSs in its upstream neighborhood, highlighted in purple box (I).

### Protein Model

*Glys* in *D. ananassae* has five CDSs within the genome sequence. The first unique protein sequence (Glys-PA) is translated from one mRNA isoform (*Glys-RA*; Figure 1B). The next unique protein sequence (Glys-PC, Glys-PB and Glys-PD) is translated from three mRNA isoforms that differ in their UTRs (*Glys-RC, Glys-RB* and *Glys-RD*; Figure 1B). Relative to the ortholog in *D. melanogaster*, the CDS number and protein isoform count are conserved, although the *BLAT* Alignments of NCBI RefSeq gene prediction for *D. ananassae* to the *D. ananassae* May 2011 (Agencourt dana_caf1/DanaCAF1) assembly incorrectly displays the first two CDSs of *Glys-RA* and the first CDS of *Glys-RC* of the putative ortholog located farther upstream (Figure 1B; Figure 1D). The sequence of Glys-PA in *D. ananassae* has 97.32% identity (E-value: 0.0) with the protein-coding isoform Glys-PA in *D. melanogaster*, as determined by *blastp* (Figure 1C). Coordinates of this curated gene models (Glys-PA, Glys-PB, Glys-PC, Glys-PD) are stored by NCBI at GenBank/BankIt (accession <u>**BK064519**, **BK064520**, **BK064521**, **BK064522**</u>, respectively). This gene model can also be seen within the target genome at this TrackHub.

### Special characteristics of the protein model

The BLAT Alignments of NCBI RefSeq Genes predicted the first two CDSs of *Glys-RA* and the first CDS of *Glys-RC* upstream of the putative ortholog, highlighted by the purple box denoted I in Figure 1D. This finding might imply a tandem gene duplication of *Glys*. However, a more current RefSeq assembly, *D. ananassae* Sep. 2021 (University of Maryland, ASM1763931v2/DanaRefSeq2), does not support the presence of the potential gene duplication and exhibits a single conserved ortholog to *Glys* in *D. melanogaster*. It is possible there was a tandem duplication of *Glys* in the strain sequenced for the assembly used in this gene annotation report (GCA_000005115.1) or that there was a misassembly inducing an apparent duplication. Given the failure to replicate the apparent duplication in the second assembly, however, leads us to the conservative conclusion that there is most likely no duplication event of *Glys* in the *D. ananassae* lineage and that there instead is likely as misassembly in this region in the focal assembly.

## Methods

“Detailed methods including algorithms, database versions, and citations for the complete annotation process can be found in Rele et al. (2023). Briefly, students use the GEP instance of the UCSC Genome Browser v.435 (https://gander.wustl.edu; Kent WJ et al., 2002; Navarro Gonzalez et al., 2021) to examine the genomic neighborhood of their reference IIS gene in the *D. melanogaster* genome assembly (Aug. 2014; BDGP Release 6 + ISO1 MT/dm6). Students then retrieve the protein sequence for the *D. melanogaster* reference gene for a given isoform and run it using *tblastn* against their target *Drosophila* species genome assembly on the NCBI BLAST server (https://blast.ncbi.nlm.nih.gov/Blast.cgi; Altschul et al., 1990) to identify potential orthologs. To validate the potential ortholog, students compare the local genomic neighborhood of their potential ortholog with the genomic neighborhood of their reference gene in *D. melanogaster*. This local synteny analysis includes at minimum the two upstream and downstream genes relative to their putative ortholog. They also explore other sets of genomic evidence using multiple alignment tracks in the Genome Browser, including BLAT alignments of RefSeq Genes, Spaln alignment of *D. melanogaster* proteins, multiple gene prediction tracks (e.g., GeMoMa, Geneid, Augustus), and modENCODE RNA-Seq from the target species. Detailed explanation of how these lines of genomic evidenced are leveraged by students in gene model development are described in Rele et al. (2023). Genomic structure information (e.g., CDSs, intron-exon number and boundaries, number of isoforms) for the *D. melanogaster* reference gene is retrieved through the Gene Record Finder (https://gander.wustl.edu/~wilson/dmelgenerecord/index.html; Rele et al., 2023). Approximate splice sites within the target gene are determined using *tblastn* using the CDSs from the *D. melanogaste*r reference gene. Coordinates of CDSs are then refined by examining aligned modENCODE RNA-Seq data, and by applying paradigms of molecular biology such as identifying canonical splice site sequences and ensuring the maintenance of an open reading frame across hypothesized splice sites. Students then confirm the biological validity of their target gene model using the Gene Model Checker (https://gander.wustl.edu/~wilson/dmelgenerecord/index.html; Rele et al., 2023), which compares the structure and translated sequence from their hypothesized target gene model against the *D. melanogaster* reference gene model. At least two independent models for a gene are generated by students under mentorship of their faculty course instructors. Those models are then reconciled by a third independent researcher mentored by the project leaders to produce the final model. Note: comparison of 5’ and 3’ UTR sequence information is not included in this GEP CURE protocol.” (Gruys et al., 2025b)

## Supplemental Files

1. Zip file containing a FASTA, PEP, GFF files for the gene model
2. Figure 1 in high resolution

## Metadata

Bioinformatics, Genomics, *Drosophila*, Genotype Data, New Finding

## Acknowledgements

This publication is dedicated to the memory of Dr. James J. Youngblom. We would like to thank Wilson Leung for developing and maintaining the technological infrastructure that was used to create this gene model. Thank you to [FlyBase](https://flybase.org/) for providing the definitive database for *Drosophila melanogaster* gene models. Further, we would like to thank the editors and developers at the journal *microPublication: Biology* for assistance in developing the template for these single gene ortholog publications.

## Funding

This material is based upon work supported by the National Science Foundation under Grant No. IUSE-1915544 to LKR and the National Institute of General Medical Sciences of the National Institutes of Health Award R25GM130517 to LKR. The Genomics Education Partnership is fully financed by Federal money. The content is solely the responsibility of the authors and does not necessarily represent the official views of the National Institutes of Health.

## Notes

https://gander.wustl.edu/cgi-bin/hgTracks?db=DanaCAF1&lastVirtModeType=default&lastVirtModeExtraState=&virtModeType=default&virtMode=0&nonVirtPosition=&position=scaffold_13340%3A6039852-6043756&hgct_customText=track%20type=bigGenePred%20visibility=pack%20bigDataUrl=http://genemodels01.ua.edu/trackhub_hosting/models/DanaCAF1/DanaCAF1.bb

### Competing Interest Statement

The authors have declared no competing interest.

### Summary of Updates

Two of the embedded hyperlinks were striped from the manuscript when PDF was created originally. The new PDF has those link live.

